# Carnivores in a metropolitan forest island: first spatial capture-recapture estimates reveal high densities of leopards and striped hyenas in Jaipur, India

**DOI:** 10.64898/2026.10.05.755896

**Authors:** Riddhi Sondagar, Kamakshi Singh Tanwar, Mohanraj Thangavel, Matt Hayward, Qamar Qureshi

## Abstract

Urban protected areas (PAs) have become important habitats for wildlife in rapidly expanding human-dominated landscapes. These PAs may provide refuge and continuously abundant resources, which may enable some adaptable carnivores to persist and attain high abundances. However, little is known about the population ecology and interactions between carnivores sharing such spatially restricted and disturbed urban habitats. We evaluated the densities and spatio-temporal interactions of leopards (*Panthera pardus*) and striped hyenas (*Hyaena hyaena*) in Jhalana-Amagarh Conservation Reserve, a protected area (PA) embedded within the metropolis of Jaipur, India. During May-July 2025, we deployed 30 paired camera-traps and used spatially explicit capture-recapture models to estimate density and individual activity centres. We further quantified their diel activity and temporal overlap using kernel density estimation. Leopard density was estimated at 7.03 ± 1.31 individuals/10 km² and striped hyena density at 4.06 ± 0.96 individuals/10 km², with activity centres of both species occurring in close proximity and frequently towards the reserve periphery. Both species were predominantly nocturnal and showed high temporal overlap (Δ^_4_=0.80, CI=0.73-0.86). Our results report exceptionally high leopard and striped hyena densities in an urban PA and exemplify the importance of small PAs within rapidly expanding cities, stressing the need for finer-scale studies more broadly to understand the mechanisms shaping interactions between the two species. However, such systems require careful management of food subsidies, habitat quality and connectivity along with long-term monitoring of population viability, disease dynamics and human-wildlife conflict.

## Introduction

Natural habitats continue to lose ground to anthropogenic infrastructure as the world becomes more urbanised (Hamer and McDonnell 2010). In such cases, wildlife can either be excluded from urban spaces or adapt to human-modified landscapes. While urbanisation presents one of the greatest threats to global biodiversity (Seto et al. 2012), cities can be crucial for conserving native biodiversity through the semi-natural and artificial ecological systems present within and around them (Ives et al. 2016; Soanes and Lentini 2019). As two-thirds of the world’s human population is projected to live in urban areas by 2050 (United Nations, 2019), human-wildlife interactions will increase and reshape our understanding of wildlife ecology in human-dominated environments (Aronson et al. 2017). Across the world, habitats in urban areas exhibit some similar ecological characteristics, even if present in very different biogeographical locations (Savard et al. 2000; Groffman et al. 2014). This novel environment presents unique challenges to wildlife, including the presence of vehicular and pedestrian traffic, light and noise pollution, and a decline of natural food resources (Grimm et al. 2008). The heat island effect of urban areas may also present higher temperatures than their surroundings (Oke 1982). While urbanisation leads to a loss of species diversity (McKinney 2006), a subset of behaviourally flexible, ecological generalists can successfully exploit human-dominated environments and often occur at relatively high numbers (termed synurbic species) (Bateman and Fleming 2012; Athreya et al. 2013).

Mammalian terrestrial carnivores appear sensitive to human development owing to their expansive home ranges, low population densities and high trophic positions (Terborgh 1974; Woodroffe 2000; Barker et al. 2023). Their potential for human-wildlife conflict also makes carnivores the first to be lost from urban landscapes (Crooks 2002; Bateman and Fleming 2012). Large carnivore avoidance of humans has been attributed either to perceived predation risk (fear) (Oriol-Cotterill et al. 2015; Smith et al. 2017) or behavioural adjustments to anthropogenic disturbance (Frid and Dill 2002). Despite barriers imposed by urban environments, several carnivores have established themselves in numerous cities worldwide. Examples include coyotes (*Canis latrans*) in Chicago, United States (Gehrt et al. 2011), caracals (*Caracal caracal*) in Cape Town, South Africa (Leighton et al. 2022), red foxes (*Vulpes vulpes*) in cities of Great Britain (Scott et al. 2014), golden jackals (*Canis aureus*) in Kolkata, India (Gupta et al. 2016) and striped hyenas (*Hyaena hyaena*) in Jaipur, India (Yosef and Kumbhojkar 2025). Leopards (*Panthera pardus*) have been documented from various cities including Nairobi, Kenya (Landy et al. 2018), as well as Mumbai, Maharashtra (Surve et al. 2022), Guwahati, Assam (Deka et al. 2025), Delhi (Chaudhary et al. 2023), and Bengaluru, Karnataka (Kolekar et al. 2024) in India. However, with the exception of domestic cats, very few felids can be considered established urban dwellers, as they are mostly confined to habitat pockets (like gardens, parks or forest patches) instead of residing in the built-up areas of the city (Bateman and Fleming 2012).

A growing body of scientific evidence suggests that the persistence of carnivores in urban landscapes is underpinned by behavioural plasticity and ecological flexibility that allow them to exploit anthropogenic environments and reduce the risks associated with human activity (Soulsbury and White 2015; Gese and Bekoff 2004, Gaynor et al. 2018). Cities provide easy, abundant and non-seasonal food sources (synanthropic prey, road-kill and human refuse) (Baker et al. 2000; Kumar et al. 2019; Reshamwala et al. 2018) as well as permanent water sources. Consequently, the survival, growth rate, body condition and population densities of urban dwellers fare better than their wilder counterparts (Fedriani et al. 2001; Webbon et al. 2004; Davison et al. 2009). Such increases in resource availability and productivity can also modify relationships between sympatric predators (Newsome et al. 2015).

Leopards are the most widespread of large carnivores. They are also the most likely large felid to coexist with humans and livestock (Jacobson et al. 2016), enabling them to persist across the full spectrum of human-modified landscapes, from agricultural mosaics to densely populated urban areas (Vijayan and Pati 2002; Devens et al. 2021; Surve et al. 2022). This close proximity of leopards with humans has also made them responsible for a large number of attacks on humans (Goyal et al. 2000; Singh 2005). Interestingly, leopards are known to provide ecosystem services in urban landscapes by suppressing populations of free-ranging dogs thereby potentially reducing the risk of dog bites and rabies transmission (Braczkowski et al. 2018; Kumbhojkar et al. 2025). Given their potential for both ecosystem benefits and human-wildlife conflicts, it is imperative we accurately estimate their population sizes in our cities.

Leopards rarely occur in isolation, even when living in the habitat pockets of urban cities (Surve et al. 2022). They coexist with other carnivores and may overlap in habitat use, activity patterns and diet, thereby creating opportunities for both competition and facilitation. The striped hyena is distributed widely across India and frequently occurs in landscapes occupied by leopards (Jhala et al. 2021). Unlike in protected areas (PAs) (Arivazhagan et al. 2007; Mandal 2018; Ashish et al. 2024; Dutta 2024), interactions between the two species outside PAs have received little scientific attention. Long-term conservation of these carnivores depends on understanding both the extent to which they can tolerate human activities and the behavioural adaptations and ecological conditions that enable their persistence in human-dominated landscapes. Such knowledge can only be gained by studying populations living in and around urban areas and is essential for designing connected landscapes in which PAs are linked through compatible multiple-use habitats.

In this paper, we estimate the population densities of leopards and striped hyenas in Jhalana-Amagarh Conservation Reserve, an urban PA in Jaipur, the tenth largest metropolitan region of India (Jawaid et al. 2017). Given the paucity of data on interactions between leopards and striped hyenas outside PAs, we also examined their patterns of spatial association and temporal overlap. We hypothesised that (a) food subsidies present near cities would support relatively high densities of both carnivores and (b) limited availability of suitable habitat would result in substantial spatial proximity between leopards and striped hyenas, with high temporal partitioning potentially facilitating their ecological separation. By examining their spatial and temporal patterns, we aim to understand how large carnivores interact within increasingly human-dominated environments.

### Study Area

Jhalana-Amagarh Conservation Reserve (hereafter JACR; Figure 1) (26.955205° - 26.833211° N and 75.820879° - 75.885315° E) was gazetted in 2023 as a PA of approximately 38 km^2^ located on the south-eastern side of Jaipur in Rajasthan, India. Historically a hunting ground for royal families, Jhalana was designated as a reserved forest in 1961, under the Rajasthan Forest Act 1953, and was subsequently declared a PA in May 2017, as India’s first leopard reserve. Amagarh was protected under state environmental laws as a reserved forest and served as a corridor and overflow habitat for Jhalana for years before becoming the second leopard reserve in 2022. Managed as a single PA today, Jhalana and Amagarh are divided by several settlements and the National Highway 11 (NH11). The entire reserve is surrounded by densely populated human settlements and is heavily encroached.

**Figure 1.**
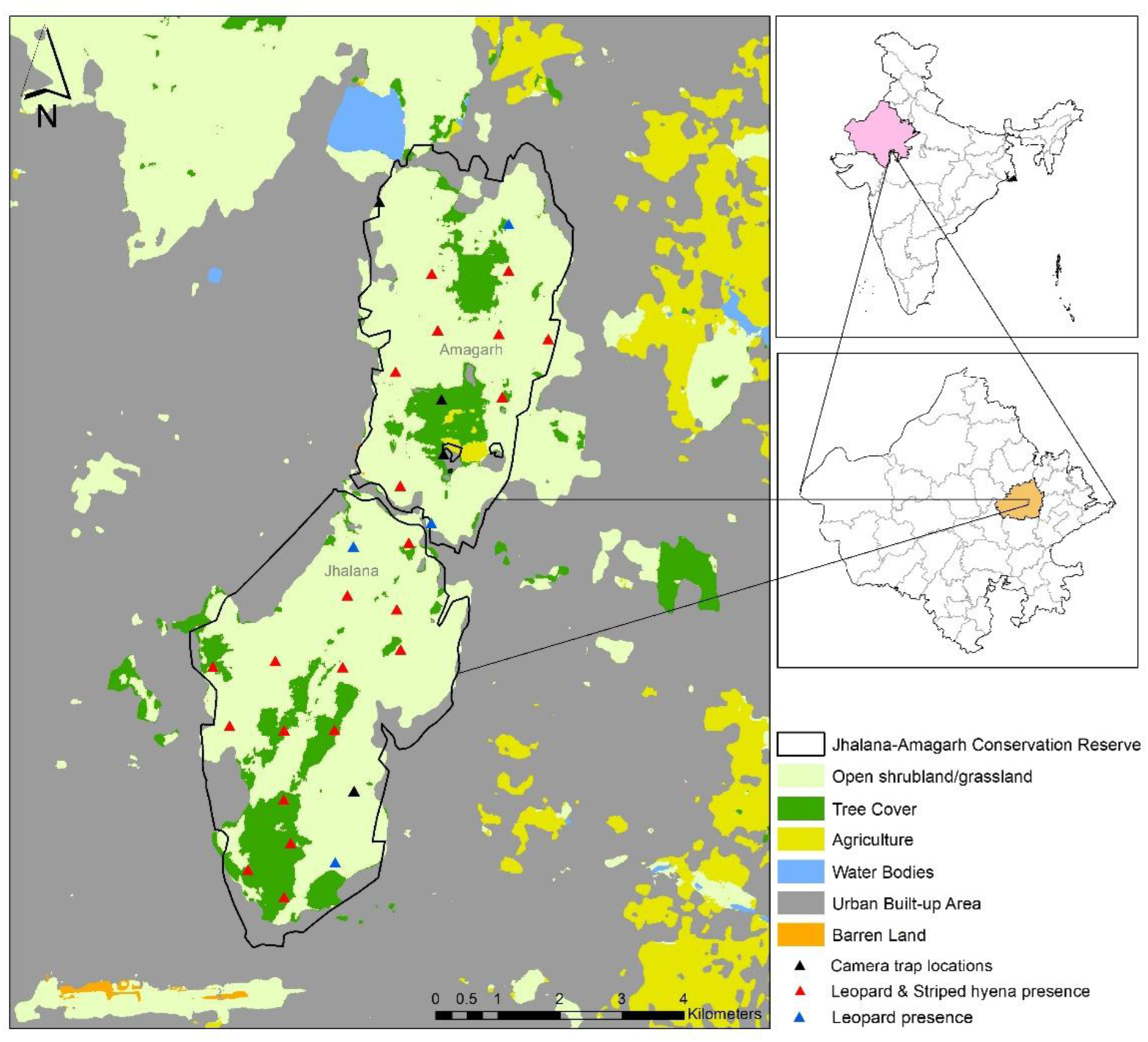
Land-use composition and spatial detections of leopards and striped hyenas (blue and red triangles) within the camera trapping array (black triangles) in Jhalana-Amagarh Conservation Reserve, India.

JACR forms a part of the Aravalli Range and hence has undulating to hilly terrain interspersed with plains and valleys. The reserve consists of northern tropical dry deciduous forest and northern tropical thorn forest (Champion and Seth 1968) where trees shed their leaves during the dry season while turning lush green after the onset of monsoon. The valley floor of the reserve is dominated by Tortilis (*Acacia tortilis*) and Khejri (*Prosopis cineraria*) while the hill slopes support a variety of native flora including Dhonk (*Anogeissus pendula*), Salar (*Boswellia serrata*), Kumta (*Acacia senegal*) and Goya kher (*Dichrostachys cinerea*). The reserve is infested with the invasive Vilayati babool (*Prosopis juliflora*) (https://forest.rajasthan.gov.in).

The climate of the study area is tropical dry with four distinct seasons: summer (March to June), monsoon (July to August), post-monsoon (September to October) and winter (November to February). The average annual precipitation in Jaipur is around 600 mm (Jaipur Development Authority 2025) with most rain recorded during the monsoon season. The reserve lacks perennial natural water sources, with only a few ephemeral streams flowing during the monsoon. Hence, artificial waterholes have been created and are maintained by the forest department to provide year-round water availability for wildlife.

The leopard is the apex predator in the reserve and occurs with a diverse carnivore guild comprising striped hyena (*Hyaena hyaena*), golden jackal (*Canis aureus*), desert fox (*Vulpes vulpes pusilla*), honey badger (*Mellivora capensis*), jungle cat (*Felis chaus*), rusty spotted cat (*Prionailurus rubiginosus*), ruddy mongoose (*Urva smithii*), grey mongoose (*Herpestes edwardsi*), small Indian civet (*Viverricula indica*) and common palm civet (*Paradoxurus hermaphroditus*) amongst others. Amongst wild herbivores, nilgai/blue bull (*Boselaphus tragocamelus*) is the most commonly visible large ungulate, with spotted deer/chital (*Axis axis*) and sambar (*Rusa unicolor*) also present, though less visible. The reserve is also home to Indian crested porcupine (*Hystrix indica*), Indian peafowl (*Pavo cristatus*), black-naped hare (*Lepus nigricollis*) and gray langur (*Semnopithecus entellus*). Domestic animals such as feral dogs (*Canis lupus familiaris*), cats (*Felis catus*), cattle (*Bos taurus*) and goats (*Capra aegagrus hircus*) are also seldom seen in the reserve.

The reserve is ringed by a 5 m tall barrier (wall + fence) (Kumbhojkar et al. 2021) that separates the forest from the high density of permanent human settlements of Jaipur city. According to the 2011 census (last census of India), Jaipur’s population is a little over 3 million, with a 10% floating population (Jaipur Municipal Corporation, http://www.jaipurmc.org) making it the 10th most populous city of India. Large quantities of uncollected refuse and carrion accumulate in the surrounding human-dominated landscape, which provides abundant food resources for a sizeable population of free-ranging dogs. Despite the presence of a boundary wall along most of the reserve, it is permeable to both people and wildlife. Local residents frequent the fringes of the reserve in search of firewood and for grazing their livestock and wild animals regularly move beyond its boundaries into the urban landscape.

## Methodology

### Camera trapping

Thirty (30) pairs of camera traps were systematically deployed in a 1 x 1 km^2^ grid covering the entire area of JACR in two blocks sequentially from 14^th^ May to 29^th^ July 2025 (Figure 1). Camera traps were strategically deployed on animal trails, forest roads and patrol routes, which apex predators preferentially use (Karanth and Nichols 1998; Jhala et al. 2011; Hayward and Marlow 2014), following information from a reconnaissance sign survey conducted prior to the study. In each grid, a pair of double-sided Cuddeback C1 cameras (*n=*60) were placed approximately 30-45 cm above ground and 3-6 m across the trails to maximize the likelihood of capturing full-body images of leopards, striped hyenas and other animals (Jhala et al. 2011). A minimum distance of 485 m was maintained between camera stations. Cameras operated continuously, day and night, with ‘fast as possible’ trigger delay, full aspect ratio, and no video or time-lapse mode. Stations were checked weekly for battery status, maintenance and data retrieval.

The two contiguous blocks, with a 15-day overlap in sampling, were pooled into a single closed spatially explicit capture-recapture (SECR) session because the same individuals were detected in both blocks, indicating a shared spatially connected population. Moreover, the 77-day sampling period fell within the commonly used 90-day guideline for closed population sampling (Karanth and Nichols 2002).

While grid-based sampling of 2 km^2^ is commonly employed for estimating populations of large carnivores such as tigers and leopards (Jhala et al. 2011), we adopted a finer grid size of 1 km^2^ (Sehgal et al. 2023). Since this was the first systematic camera-trap study in the reserve, a finer spatial resolution allowed comprehensive coverage of the study area and provided an opportunity to capture information on small-ranging and elusive species in addition to large carnivores.

### Analytical Methods

#### Individual identification of leopards and striped hyenas

All camera trap records were manually processed and identified to species level. In species with natural pelage markings, like leopards and striped hyenas, the rosettes and stripes on them are prominent and unique to each individual. We identified individual leopards and striped hyenas by these patterns present on both their right and left flanks and limbs. Because most large carnivores are sexually dimorphic, differences in their sex-based morphological traits (Balme et al. 2012; Gupta et al. 2009; Hiby et al. 2009; Jhala et al. 2021) further helped identify individuals and apply complex heterogeneity models for estimating their densities. We first matched images of a species flank-wise and later merged them to form a single individual based on the time of each image of both flanks captured from either side of the camera station. The individuals thus identified were given individual IDs. Since complementary captures for both the species were not available for all individuals, for analysis we retained the flank having the highest number of identifiable individuals which was the left flank (30 leopards, 19 striped hyenas), in addition to the individuals photographed on both sides, while excluding unclear images and uncertain individuals.

#### Abundance and density

To estimate densities of leopards and striped hyenas in our study area, we employed the maximum-likelihood based SECR method (Borchers and Efford 2008; Kalle et al. 2011; Mandal 2018). We developed a trap matrix for each camera station and detection histories for each identified individual, considering one occasion as a 24-hour period. We then used the functions ‘secr.fit()’ and ‘region N’ within the ‘secr’ package (version 5.4.3; Efford 2026) in R (version 4.6.0; R Core Team 2026) to estimate densities and abundance of leopards and striped hyenas using various habitat-buffer masks.

In SECR, habitat-buffer mask represents the area around the camera trap array that the animals can potentially occupy and is typically defined as multiples of the suggested movement parameter (*σ*, sigma) of the study species. Owing to the unique landscape of our study area bordering dense urban areas of Jaipur city and considering the ecology of our study species, we estimated their density in experimental habitat masks with varying buffer widths. The initial *σ* for leopard and striped hyena in the study area were 607.5 and 746.4 m respectively, hence we increased the buffer widths based on these values. Although a buffer of over 4 × *σ* is commonly recommended (Efford 2025), the unavailability of suitable habitat beyond the PA necessitated the use of conservative buffer widths of 700 m, 1000 m and 2000 m from the PA boundary. Density estimates were compared across these buffer widths to evaluate their sensitivity to the assumed extent of available habitat and to determine buffer adequacy based on the stability of density estimates with increasing buffer widths, following the principle of buffer independence (Efford et al. 2022). We assumed that species movement outside the PA was restricted by human habitation and therefore developed our masks to primarily include agricultural fields and small vegetation patches while excluding urban built-up areas.

Furthermore, to ensure that the SECR model yielded stable and reliable estimates, convergence of likelihood and densities were assessed by refitting the selected SECR model using multiple sets of substantially different starting values. The resulting density (*D*) and movement-scale (*σ*) estimates showed no to negligible variation among runs. Therefore, it was concluded that the high consistency of density (*D*) and movement-scale (*σ*) estimates across runs, together with convergence of the likelihood optimization, indicated stability of the parameter estimates and robustness of the density estimate to the choice of starting values.

Using the Half-Normal (HN) function (Efford 2004), a total of seven different SECR models were fitted in all three habitat-buffer widths. The heterogeneity models for the leopard density included the effect of known sex on baseline detection probability (*g*0) and movement parameter (*σ*), while for striped hyena, owning to the inability to identify the sex, two-class finite mixture models were used where the classes are considered as a latent variable. Density (*D*) was modelled as constant for all models. A 100-m spacing was used to provide sufficiently fine spatial resolution for representing the SECR habitat mask relative to the spatial scale of detection indicated by the preliminary *σ* estimates. This resolution provided adequate representation of the detection process without unnecessarily increasing computational demand.

#### Activity centres and spatial association

Individual activity centres were estimated from the best fitted SECR model using the function ‘fxiMode()’ in the ‘*secr’* package (Efford 2004; Efford 2026; Borchers and Efford 2008). These estimated coordinates represent model-based (maximum-likelihood) modal activity centre for each individual and are subject to estimation uncertainty. The coordinates were then exported to ArcMap version 10.8, where they were used to visualise distribution and analyse spatial association. We calculated the inter and intraspecific distances between the activity centres using the ‘generate near table’ tool. Spatial association or proximity between individuals and both species was assessed by manually comparing the relative nearness of their modal activity centres.

#### Activity pattern and temporal overlap

To evaluate the temporal segregation between leopards and striped hyenas in our study area we followed two steps. First, we assessed their diel activity pattern using the non-parametric kernel density estimation (KDE) function in the ‘*overlap’* package (Ridout and Linkie 2009; Meredith and Ridout 2014) in R (version 4.6.0). KDE estimates the probability density over a 24 h cycle, where the area under the curve equals one, representing the relative distribution of activity throughout the day. We used independent captures separated by 30-minute intervals (O’Brien et al. 2003) to analyse the temporal patterns of both species.

Second, we measured the extent of overlap between the diel activity pattern of both species using the overlap coefficient (Δ), which ranges from 0 (complete segregation and no overlap) to 1 (complete overlap). The overlap was measured on the basis of ‘Dhat or Delta’ where a value of >0.75 signifies a high overlap following Monterroso et al. (2014). Since our sample size was greater than 75, we calculated the Delta^4^/dhat4 (Δ^_4_) values (Meredith et al. 2013; Ridout and Linkie 2009). The precision of the overlap coefficient (Δ) was assessed by calculating the 95% confidence interval (95% CI) using 2000 bootstrap resampling iterations (Ridout and Linkie 2009).

Additionally, we calculated the percent time active during the day for both species using the function ‘fitact()’ in ‘*activity’* package (Rowcliffe and Rowcliffe 2016), in R.

## Results

### Sampling effort

Camera traps were operable for between 20 and 77 days giving a total trapping effort of 1111 trap nights. Relatively short periods at some stations arose from limited camera availability, camera failures and disturbance by animals. The segregated data from camera trapping yielded a total of 389 leopard and 550 striped hyena captures, of which 365 and 410 images respectively were suitable for individual recognition. We identified a total of 30 leopard individuals, of which 17 were females, 9 were males and 4 were of unidentifiable sex, along with 19 striped hyena individuals. Out of 30 camera trap locations, leopards and striped hyenas were captured in 26 and 22 locations respectively (Figure 1).

#### Abundance and density

To assess the sensitivity of density estimates and associated baseline parameters for the two species to the extent of the state-space mask, we compared the results across multiple buffer widths (700, 1000, 2000 m) and found the estimates remained largely consistent across them (Table 1). Density estimates from various buffers widths exhibited small differences in coefficients of variance and substantially overlapping 95% confidence intervals (Table 1). We therefore retained the 1000 m buffer for inference as further increases in buffer width produced negligible changes in the maximized likelihood, density estimates, and associated parameters. Given the limited availability of suitable habitat beyond the reserve boundary, extending the state-space mask further into the surrounding urban areas was also considered biologically unrealistic.

**Table 1.** Density and baseline parameter estimates of leopard and striped hyena across various buffer widths surrounding Jhalana-Amagarh Conservation Reserve, India.

| Species | Buffer width (meters) | Model | Density /10 km<sup>2</sup> | Density SE | Density LCL | Density UCL | g0 | g0 SE | sigma | sigma SE | pmix | pmix SE |
| --- | --- | --- | --- | --- | --- | --- | --- | --- | --- | --- | --- | --- |
| Leopard | 700 m | g0sh2 (Female) | 7.34 | 1.36 | 5.11 | 10.54 | 0.17 | 0.01 | 420.83 | 28.12 | 0.74 | 0.07 |
|  |  | g0sh2 (Male) | 7.34 | 1.36 | 5.11 | 10.54 | 0.17 | 0.01 | 765.61 | 47.55 | 0.25 | 0.07 |
|  | 1000 m | g0sh2 (Female) | 7.03 | 1.31 | 4.89 | 10.12 | 0.18 | 0.02 | 418.99 | 28.64 | 0.75 | 0.07 |
|  |  | g0sh2 (Male) | 7.03 | 1.31 | 4.89 | 10.12 | 0.18 | 0.02 | 807.49 | 52.11 | 0.24 | 0.07 |
|  | 2000 m | g0sh2 (Female) | 6.84 | 1.28 | 4.75 | 9.85 | 0.18 | 0.02 | 417.60 | 27.99 | 0.76 | 0.07 |
|  |  | g0sh2 (Male) | 6.84 | 1.28 | 4.75 | 9.85 | 0.18 | 0.02 | 844.42 | 60.82 | 0.23 | 0.07 |
| Striped hyena | 700 m | g0sh2 (sex undefined) | 4.37 | 1.02 | 2.77 | 6.87 | 0.21 | 0.01 | 855.44 | 44.22 | 0.43 | 0.12 |
|  |  |  | 4.37 | 1.02 | 2.77 | 6.87 | 0.21 | 0.01 | 438.58 | 28.00 | 0.56 | 0.12 |
|  | 1000 m |  | 4.06 | 0.96 | 2.57 | 6.42 | 0.21 | 0.01 | 852.22 | 45.21 | 0.45 | 0.13 |
|  |  |  | 4.06 | 0.96 | 2.57 | 6.42 | 0.21 | 0.01 | 439.85 | 27.83 | 0.54 | 0.13 |
|  | 2000 m |  | 3.85 | 0.91 | 2.43 | 6.09 | 0.21 | 0.01 | 441.56 | 27.60 | 0.53 | 0.13 |
|  |  |  | 3.85 | 0.91 | 2.43 | 6.09 | 0.21 | 0.01 | 852.91 | 40.67 | 0.46 | 0.13 |

**g0sh2:** $\hat{D} \sim 1$ , $\hat{g0} \sim 1$ , $\hat{\sigma} \sim \text{sex/latent classes}$ ; **SE:** Standard error; **LCL:** Lower Confidence Limit; **UCL:** Upper Confidence Limit; **g0:** Baseline detection probability at the activity centre ( $g_0$ ); **s:** Spatial scale of detection ( $\sigma$ ); **pmix:** mixture proportion.

The maximum-likelihood SECR based density of leopards in our study area was estimated at 7.03 ± 1.31 individuals/10 km^2^ (95% CI 4.89-10.12) with a constant baseline detection probability (*g*0) and estimated movement parameter (*σ*) of 418.99 m ± 28.64 SE for females and 807.49 m ± 52.11 SE for males for the best fit model (*g*0*sh*2: *D*^∼1, *g*^0∼1, *σ*^∼sex) based on the AIC (607.34; Table 2) (Table 1 and 3). As our study area is relatively small (∼ 38 km^2^), we used a 10 km^2^ unit to represent the density.

**Table 2.** Comparative evaluation of heterogeneous and non-heterogeneous spatially-explicit capture-recapture models (chosen model in bold) and their relative support for estimating leopard and striped hyena density, with comparisons conducted separately within each model class, in Jhalana-Amagarh Conservation Reserve, India.

| Site + Buffer | Species | Model Class | Model Specification | Detection Function | *npar* | AIC | ΔAIC | AIC Weight |
| --- | --- | --- | --- | --- | --- | --- | --- | --- |
| Jhalana-Amagarh Conservation Reserve + 1000 m buffer around it | Leopard | Heterogeneity models | **$g0sh2: D \sim 1 g0 \sim 1 \sigma \sim sex \text{ pmix} \sim sex$** | Half-normal | **5** | **607.34** | **0** | **0.7265** |
| | | | $gh2sh2: D \sim 1 g0 \sim sex \sigma \sim sex \text{ pmix} \sim sex$ | | 6 | 609.30 | 1.954 | 0.2735 |
| | | | $gh2s0: D \sim 1 g0 \sim sex \sigma \sim 1 \text{ pmix} \sim sex$ | | 5 | 647.40 | 40.058 | 0 |
| | | Non-heterogeneity models | $g0s0: D \sim 1 g0 \sim 1 \sigma \sim 1$ | | 3 | 614.03 | 0 | 0.5344 |
| | | | $gbs0: D \sim 1 g0 \sim b \sigma \sim 1$ | | 4 | 616.03 | 2 | 0.1966 |
| | | | $g0sb: D \sim 1 g0 \sim 1 \sigma \sim b$ | | 4 | 616.03 | 2 | 0.1966 |
| | | | $gbsb: D \sim 1 g0 \sim b \sigma \sim b$ | | 5 | 618.03 | 4 | 0.0723 |
| Jhalana-Amagarh Conservation Reserve + 1000 m buffer around it | Striped hyena | Heterogeneity models | **$g0sh2: D \sim 1 g0 \sim 1 \sigma \sim h2 \text{ pmix} \sim h2$** | Half-normal | **5** | **608.86** | **0** | **0.5908** |
| | | | $gh2sh2: D \sim 1 g0 \sim h2 \sigma \sim h2 \text{ pmix} \sim h2$ | | 6 | 609.94 | 1.078 | 0.3446 |
| | | | $gh2s0: D \sim 1 g0 \sim h2 \sigma \sim 1 \text{ pmix} \sim h2$ | | 5 | 613.29 | 4.429 | 0.0645 |
| | | Non-heterogeneity models | $g0s0: D \sim 1 g0 \sim 1 \sigma \sim 1$ | | 3 | 640.566 | 0 | 0.5344 |
| | | | $gbs0: D \sim 1 g0 \sim b \sigma \sim 1$ | | 4 | 642.566 | 2 | 0.1966 |
| | | | $g0sb: D \sim 1 g0 \sim 1 \sigma \sim b$ | | 4 | 642.566 | 2 | 0.1966 |
| | | | $gbsb: D \sim 1 g0 \sim b \sigma \sim b$ | | 5 | 644.566 | 4 | 0.0723 |

**$g0$** : Baseline detection probability at the activity centre ( $g_0$ ); **s**: Spatial scale of detection ( $\sigma$ ); **D**: Density; ***npar***: Number of parameters estimated; **h2**: Heterogeneity classes (leopard: sex and striped hyena: latent); **pmix**: Mixture proportions; **b**: Behavioural response; **AIC**: Akaike Information Criterion.

For striped hyenas, since sex could not be identified for most individuals, the heterogeneity in movement (*σ*) and detection probability (*g*0) was modelled using the two-class finite mixture (*pmix*) models. Amongst them, the best fit model (*g*0*sh*2: *D*^∼1, *g*^0∼1, *σ*^∼h2) based on the AIC (608.86; Table 2), estimated striped hyena density at 4.06 ± 0.96 individuals/10 km^2^ (95% CI 2.57-6.42) with an estimated movement of 852.22 m ± 45.21 SE and 439.85 m ± 27.83 SE for the two latent classes, reflecting heterogeneity in movement amongst individuals (Table 1 and 3). Based on the best fit model, abundance was estimated at 39 ± 4 and 22 ± 2 SE individuals in 91.35 km^2^ area for leopards and striped hyenas respectively (Table 3).

**Table 3.** Density estimates of leopards and striped hyenas in Jhalana-Amagarh Conservation Reserve, India.

| Species | Best fit model | No. of Camera traps | Trap nights | mt+1 | Density/10 km<sup>2</sup> (SE) | 95% CI | Abundance (SE) |
| --- | --- | --- | --- | --- | --- | --- | --- |
| Leopard | $g0sh2: \hat{D} \sim 1, \hat{g0} \sim 1, \hat{\sigma} \sim \text{sex}$ | 30 | 1111 | 30 | 7.03 (1.31) | 4.89-10.12 | 39 (4) |
| Striped hyena | $g0sh2: \hat{D} \sim 1, \hat{g0} \sim 1, \hat{\sigma} \sim h2$ | 30 | 1111 | 19 | 4.06 (0.96) | 2.57-6.42 | 22 (2) |

**$g0$** : Baseline detection probability at the activity centre ( $g_0$ ); **$s$** : Spatial scale of detection ( $\sigma$ ); **$D$** : Density; **$h2$** : Heterogeneity classes (leopard: sex and striped hyena: latent); **$m(t+1)$** : minimum number of individuals photo-captured; **$SE$** : standard error; **$CI$** : Confidence Interval; **$Abundance (SE)$** : Population estimates with their respective standard errors.

#### Activity centres and spatial association

Activity centres of close to 50% of the leopard and many striped hyena individuals were located close to the boundary of our study area. The activity centres of most female leopards (*n* = 17) were concentrated towards the centre of the study area while most male activity centres were near the study area boundary. The activity centres of both the species in Jhalana were distributed mainly in the centre and western side of the study area, while in Amagarh, the activity centres of both species were concentrated towards the central-eastern side of the park adjacent to the few agricultural fields present outside the boundary (Figure 2). The average distances between the activity centres were computed across six categories of inter- and intraspecies spatial separation (Table 4).

**Figure 2.**
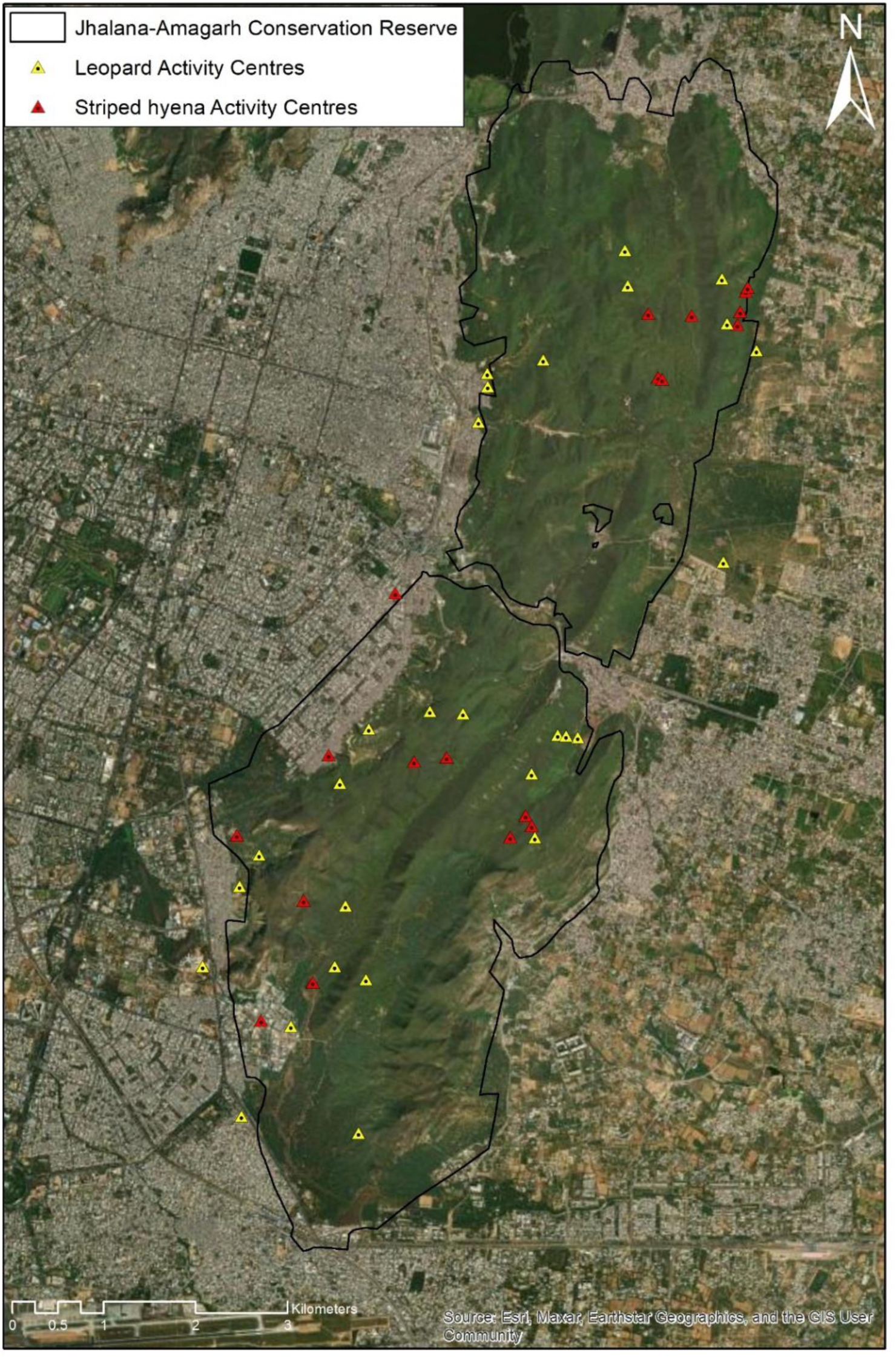
Estimated activity centre locations of leopard and striped hyena derived from best fit spatially-explicit capture-recapture model, in Jhalana-Amagarh Conservation Reserve, India, overlaid on Esri World Imagery in ArcMap version 10.8.

**Table 4.** Pairwise inter- and intraspecific distances between activity centres of leopards and striped hyenas in Jhalana-Amagarh Conservation Reserve, India.

| Distance category | No. of individuals analysis | Average Distance (km) ± SD | Minimum Distance (km) | Median (km) | Maximum Distance (km) |
| --- | --- | --- | --- | --- | --- |
| Leopard + Striped hyena | 30 leopards and 19 striped hyenas | 4.14 ± 2.63 | 0.11 | 3.90 | 10.56 |
| Leopard | 30 leopards | 4.23 ± 2.51 | 0.00 | 3.89 | 10.51 |
| Striped hyena | 19 striped hyenas | 4.19 ± 2.72 | 0.04 | 4.41 | 9.57 |
| Female Leopard | 17 females | 4.12 ± 2.41 | 0.09 | 3.82 | 10.51 |
| Male Leopard | 9 males | 4.39 ± 2.47 | 0.38 | 4.16 | 8.42 |
| Female + Male leopard | 17 females and 9 males | 4.34 ± 2.64 | 0.00 | 4.12 | 10.31 |

The unique pairwise distances between activity centres were broadly similar across species and sex categories, with averages ranging from 4.12 ± 2.41 SD to 4.39 ± 2.47 SD km (Table 4). Maximum distance was reported between leopards and striped hyena while minimum distance was reported amongst female and male leopard, and all leopard individuals (Table 4). Average distance between male leopards was greater than that between female leopards while median distances amongst all the categories were comparable, ranging from 3.82 to 4.41 km (Table 4).

#### Activity pattern and temporal overlap

A total of 208 and 246 independent captures of leopards and striped hyenas respectively were obtained using a 30-minute interval. The diel activity of leopards and striped hyenas exhibited a predominantly nocturnal pattern, with leopards extending some activity during the day and striped hyenas showing negligible activity between 09:00-16:00 IST (Figure 3). Leopards showed a slightly bimodal (nocturno-crepuscular) activity pattern with peaks between 05:00-06:00 IST and 18:30-20:00 IST while striped hyena activity was mostly concentrated between evening and early morning with the highest peak occurring between 20:00-23:00 IST at night (Figure 3). The coefficient of overlap between leopards and striped hyenas was Δ^_4_ = 0.80 (95% CI = 0.73 - 0.86). With 95% CI along the mean activity level, daily estimated activity of leopards was 58.3% (SE = 6.0%; CI = 42.4-65.8%) and that of striped hyenas was 45.0% (SE=3.0%; CI = 35.8-47.7%).

**Figure 3.**
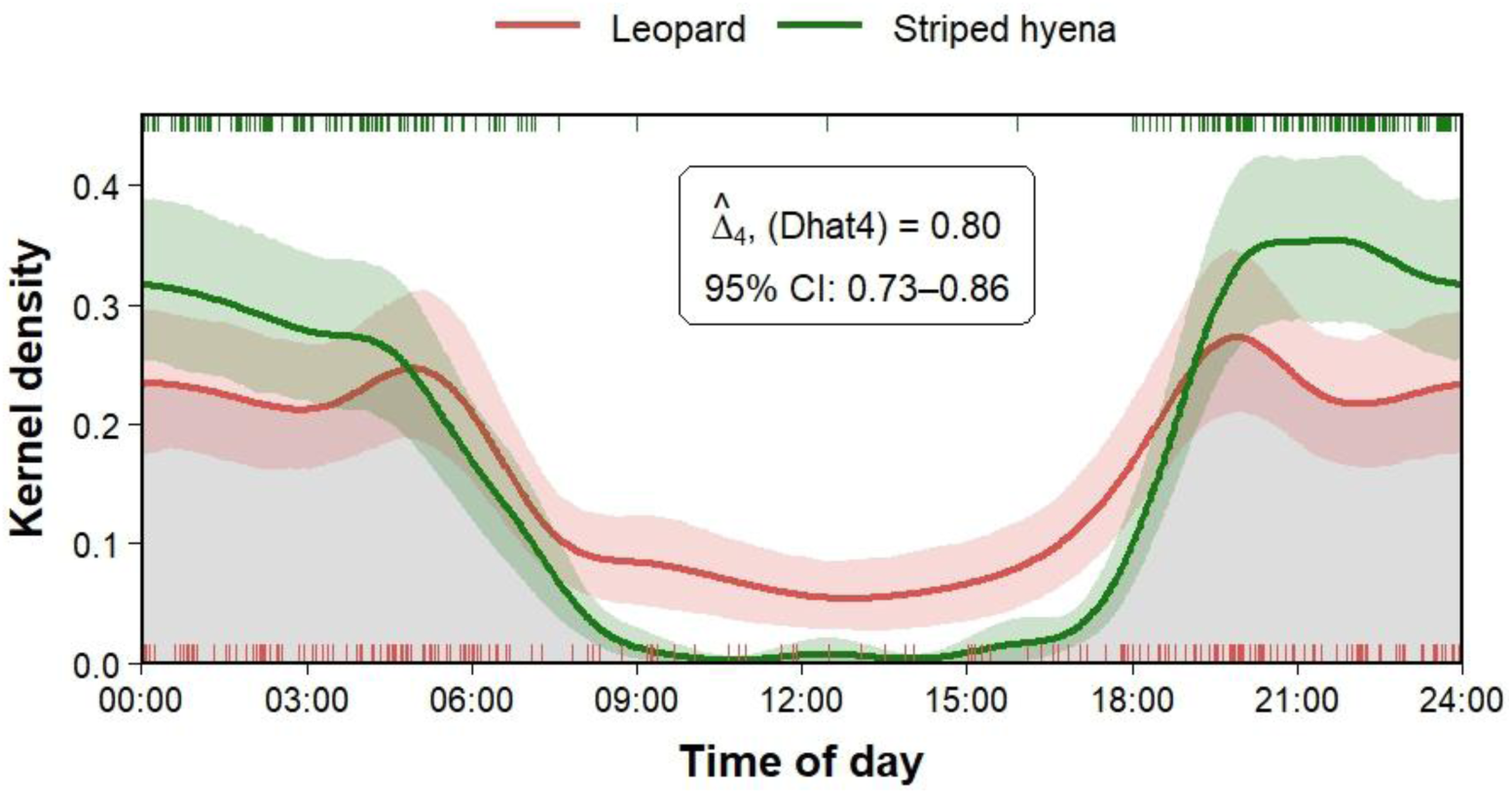
Patterns of diel activity and temporal overlap between leopards and striped hyenas in Jhalana-Amagarh Conservation Reserve, India.

## Discussion

Large carnivore conservation in human dominated landscapes comes with varying challenges. The importance of urban PAs in safeguarding wildlife, including endangered species, has been demonstrated by multiple studies conducted within India (Chaudhary et al. 2023; Surve et al. 2022; Sharma et al. 2016; Chand et al. 2025). As natural habitat continues to shrink globally, small pockets nestled within built-up areas become imperative to study and safeguard. This study provides the first SECR-based density estimates of the leopards and striped hyenas in the JACR, an island of habitat within the highly populous Jaipur city.

Leopard density recorded within our study area (70.36 ± 13.16 individuals/100 km^2^, CV-18.63%) represents one of the highest reported densities of leopards not only within India, but globally. This substantially exceeds the previous benchmark of 54.61 leopards/100 km^2^ (CV-53.04%) (Wattegedera et al. 2022) estimated by simply dividing the minimum identifiable population present at one point in time by the Kumana National Park area in Sri Lanka. Kumbhojkar et al. (2020) mentioned a higher density of 86.2 individuals/100 km^2^ from Jhalana Reserve Forest (part of our study area) which was, however, derived by dividing the total number of leopards identified over a period of two years by the administrative area of the reserve. Pooling individuals detected over an extended period and not accounting for imperfect detection or individual movement beyond the administrative boundary could have inflated the apparent density. The highest scientifically robust leopard density previously documented from India is 26.34 ± 4.96 leopards/100 km^2^ (CV-18.83%; buffer width = 8km) from Sanjay Gandhi National Park, another urban PA surrounded by the densely populated metropolis of Mumbai (Surve et al. 2022). The striped hyena density (40.69 ± 9.61 individuals/100 km²; CV-23.61%) recorded in our study was comparable to the highest reported density across the species’ range of 40.62 ± 7.36 individuals/100 km² (CV-18.12%) from Mukundara Hills Tiger Reserve (Latafat et al. 2023). Other PAs in India with high striped hyena density include Sariska (12.7 ± 2.8 individuals/100 km²; Gupta et al. 2009) and Ranthambhore Tiger Reserve (9.39 ± 1.30 individuals/100 km^2^; Latafat et al. 2023). However, our density estimates are likely an artefact of a small area acting as a refuge and might reflect habitat compression.

Notably, tiger reserves in India, which afford the highest level of protection to resident species and are usually located away from densely populated areas, thereby exhibiting low levels of anthropogenic disturbance, have reported comparatively lower leopard densities. Tiger reserves having high leopard densities include Sariska (21.43 ± 1.91 individuals/100 km²), Ranipur (16.93 ± 2.21 individuals/100 km²), Panna (16.09 ± 1.05 individuals/100 km²) and Anamalai (15.32 ± 2.99 individuals/100 km²) (Qureshi et al. 2023). Highly adaptable carnivores have been seen to exploit the proximity to human populations to their benefit and grow in number (Baker and Harris 2004; Yirga et al. 2013; Šálek et al. 2015; Panda et al. 2022; Panda et al. 2023). Leopards, perhaps more than any other large carnivore, persist in close proximity to people, including in densely settled urban and agricultural landscapes (Athreya et al. 2013; Majgaonkar et al. 2019; Surve et al. 2022). Likewise, scavengers like striped hyenas can thrive in areas where habitat is available for refuge in the daytime and carrion is available for food, possibly from the neighbouring human-dominated areas (Latafat et al. 2023). This could mean that these species have been successful in their process to adapt their behaviour and resource use to the proliferating urban environments.

Within India, leopards and striped hyenas are widely distributed beyond PAs and their occurrence is often associated with the availability of domestic prey (Majgaonkar et al. 2019). Among the domestic prey, leopards’ penchant for free-ranging dogs, that are readily available in Indian cities, could be attributed to their high densities or relative scarcity and accessibility of other prey, as they are not preferentially taken (Hayward et al. 2006, 2022). This proclivity of leopards towards dogs, as prey, has been documented throughout India in shared landscapes (Athreya et al. 2016; Ahmed et al. 2022; Desai et al. 2021) as well as in PAs located close to human habitations (Shah et al. 2009; Tiwari 2008; Surve 2015). Kumbhojkar et al. (2020) in Jhalana forest reserve found that 89.5% of leopard diet comprised domestic animals of which 44% were dogs, 16% goats and 15% cattle. Striped hyenas also exhibit considerable dietary breadth that usually corresponds to local food availability (Chourasia et al. 2012; Bhandari et al. 2020; Alam and Khan 2015; Panda et al. 2023). They are also opportunistic kleptoparasites with instances of their scavenging or appropriating kills of sympatric carnivores seen across their range (Périquet et al. 2015) including our study area (Yosef and Kumbhojkar 2025). Hence, we hypothesise that the high density of leopards in our study area is supported by the easily accessible domestic prey in the surrounding Jaipur city (Hiby et al. 2011) and refuge provided by the JACR. We supported this interpretation with the anecdotal observations of the management staff who reported that leopards frequently venture beyond the reserve boundary to prey on domestic animals and scavenge refuse from nearby meat shops. Similarly, the high density of striped hyenas may be supported by their broad and opportunistic diet which allows them to exploit anthropogenic food resources through scavenging and kleptoparasitism, while using the conservation reserve as refuge. This could also explain why the activity centres of most leopards and striped hyenas were largely concentrated towards the periphery of the study area.

Large carnivores defend territories to safeguard prey and reserve exclusive access to mates (Macdonald 1983). Across species, their home range sizes are directly proportional to body mass (McNab 1963; Lindstedt et al. 1986) and inversely proportional to resource availability (Carbone and Gittleman 2002; Hayward et al. 2009). If prey is present in abundance, carnivore territories shrink (Karanth et al. 2004) even in shared landscapes (Odden et al. 2014). Since leopards in our study area primarily depend on domestic prey present beyond the park boundary (Kumbhojkar et al. 2021), it might not make sense for them to defend large territories and expend energy. The relatively short distances between the activity centres of conspecifics as well as between leopards and striped hyenas in our study area are also perhaps understandable given the high abundance of both species and the paucity of available habitat. Additionally, by virtue of propinquity to leopard activity centres, striped hyena may get access to increased scavenging opportunities, as they are known to exploit kills made or used by sympatric carnivores as well as feed alongside leopards at the same carcass (Mandal 2018). Thus, the proximity of leopard and striped hyena activity centres might reflect both the spatial concentration of two exceptionally abundant carnivore populations within a restricted habitat and their exploitation of concentrated food resources within and around the reserve.

As human presence increasingly permeates wildlife habitats, animals have increased their nocturnality as a temporal refuge from human disturbance (Gaynor et al. 2018). Activity pattern of leopards and striped hyenas in our study area was predominantly nocturnal, with a high degree of temporal overlap. This shared shift towards nocturnality could perhaps be a common response to high human disturbance during the day, wherein the need to avoid humans outweighed the need to avoid each other (Jhala et al. 2021; Searle et al. 2021; Dahya et al. 2023; Panda et al. 2022). Patra et al. (2026) obtained a similar pattern from a shared landscape in terai, north India where leopards and striped hyenas were found largely nocturnal showing a high temporal overlap; overlap increased in areas experiencing greater human disturbance. Although both species are generally considered nocturnal, their activity often extends into day-time hours in PAs with relatively low human disturbance (Karanth et al. 2017; Tanwar et al. 2021; Dutta 2024).

In this study, we have shown that leopards and striped hyenas occur at a very high density in such a small PA and exhibit minimal spatio-temporal partitioning to mitigate potential interactions, according to metrics studied by us. Whether this segregation is sufficient to maintain reproductive success and long-term population viability remains unknown. Additionally, it may be possible that due to high resource availability, the potential for interference competition between leopards and striped hyenas is reduced (Miller et al. 2018; Chatterjee et al. 2023) or maybe hyena benefits from the high spatiotemporal overlap with leopard (Vissia et al. 2022) as that may provide striped hyenas with increased opportunities to exploit leopard-associated food resources, including scavenging from leopard kills (Mandal et al. 2018). Nevertheless, the processes operating in these populations warrant a deeper and long-term investigation.

Our results also accentuate the conservation importance of relatively small habitat patches within rapidly expanding cities, but their value will ultimately depend on how they and the landscape around them is managed. The northern extent of our study area is connected to a network of PAs, including Nahargarh Wildlife Sanctuary, its large eco-sensitive zone and Jamwaramgarh Wildlife Sanctuary, interspersed with some unprotected forest patches and sporadic agricultural fields. These habitat patches may act as stepping stones ultimately leading to Sariska Tiger Reserve (Figure 4) and may provide opportunities for the high-density leopard and striped hyena populations to move northward of the study area to occupy new territory. Retaining these habitat patches and preventing further fragmentation should therefore be a management priority, particularly as ongoing urban expansion continues to constrain habitat connectivity. Future research should assess the functional connectivity of this network using movement, occupancy or genetic approaches to determine whether leopards and other wildlife actively use these patches for movement between our study area and larger PAs like Sariska Tiger Reserve. The exceptionally high densities recorded in our study area should therefore not be interpreted as a successful result of habitat conservation or evidence that these populations require little management. At the same time, the apparent use of anthropogenic food resources presents a complex management and ecological problem. While such resources may presently contribute to high abundance of carnivores, they can also lead to increased lethal encounters, disease transmission, altered predator-prey relationships and changes in the structure and functioning of ecological communities (Newsome et al. 2015; Lanszki et al. 2018). Over the past several years, media reports have documented repeated leopard incursions into the residential areas, schools, hotels and factories in and around Jaipur city, which have occasionally resulted in attacks on people and instances of retaliatory killing of leopards. Such incidents indicate that increasing encounters at the urban interface may pose a risk to human safety and public tolerance of carnivores. Hence, efforts should be made to understand the impact of such high carnivore densities, the role of anthropogenic food subsidies and closely monitoring how changes in their availability influence the ecology of the wildlife and their tendency to move out of the reserve boundary. Finally, management should favour proactive measures that reduce opportunities for conflict in the future and foster coexistence.

**Figure 4.**
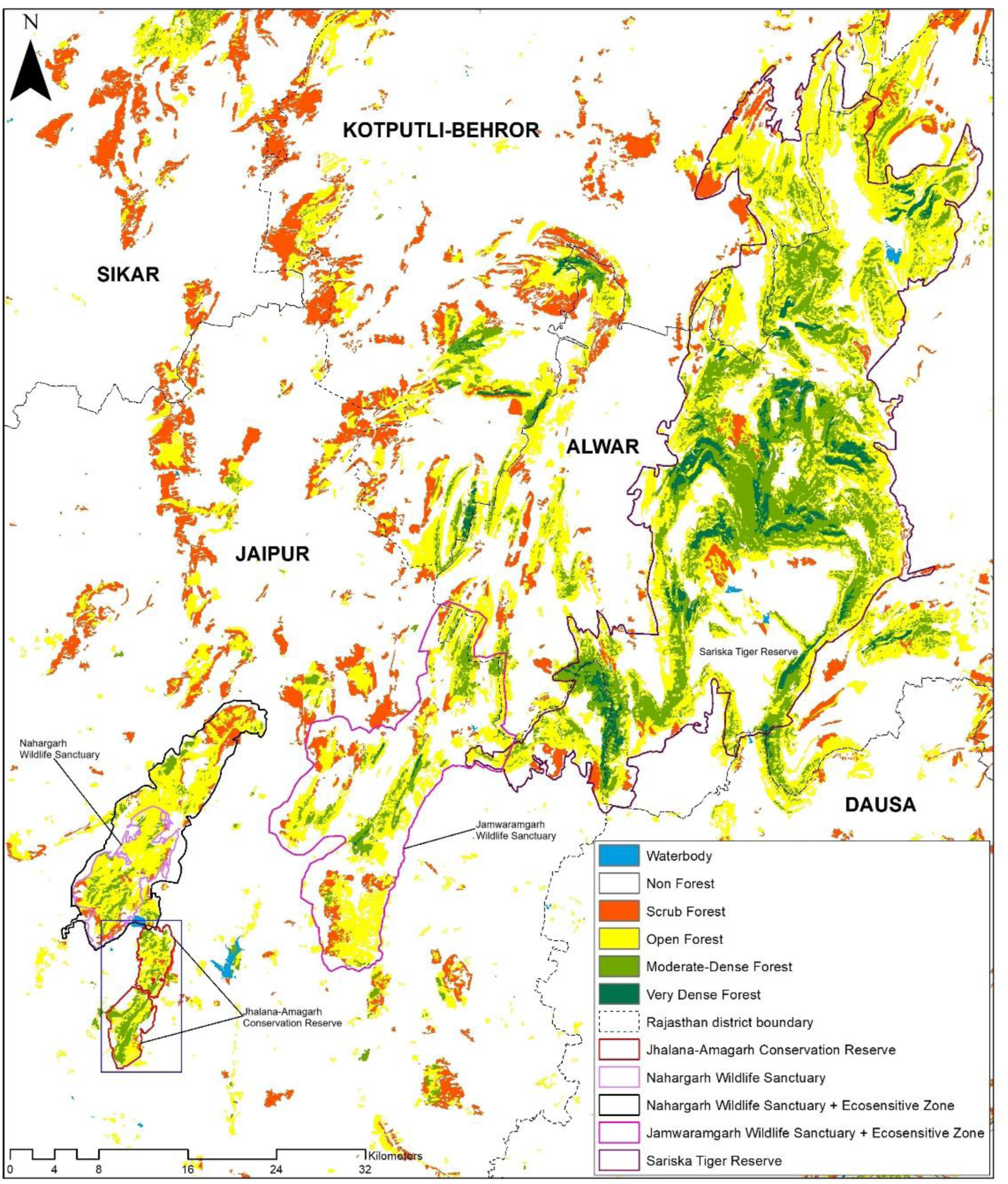
Potential landscape linkages between Jhalana-Amagarh Conservation Reserve (rectangle box) and the greater Sariska Tiger Reserve landscape (Forest-cover data: Forest Survey of India 2019; Protected Area boundaries: official website of the Forest Department, Government of Rajasthan).

## Availability of data

The data used for the study can be made available upon reasonable request to the corresponding author.

## Authors’ contributions

RS and QQ conceived the study. RS did the field work and data processing. RS and KST performed the analysis and drafted the original manuscript. MT secured the necessary permissions and logistics for the field work. RS, KST, QQ and MH interpreted the results. All authors reviewed and approved the final manuscript.

## Acknowledgements

We thank the Principal Chief Conservator of Forests and Chief Wildlife Warden, Rajasthan Forest Department for necessary permissions to carry out the field work. We extend our gratitude to the Deputy Conservator of Forests and Range Officer of Jhalana-Amagarh Conservation Reserve for arranging the logistics during the study period. Heartfelt thanks to Vikas Verma for his help with the data processing. Sincere thanks to Nishant Bhardwaj for his support and assistance during the field work. The field staff and drivers of the reserve are fondly acknowledged for their efforts and contributions during the survey.

## References

Ahmed, K., Zehra, N., & Khan, J. A. (2022). Leopard (Panthera pardus) Food habits in shared landscapes: a case study of two watersheds. International Journal of Ecology and Environmental Sciences, 48(6), 729–736.

Alam, M.S., & Khan, J.A. (2015). Food habits of striped hyena (Hyaena hyaena) in a semi-arid conservation area of India. J. Arid Land, 7(6), 860–866.

Arivazhagan, C., Arumugam, R., & Thiyagesan, K. (2007). Food habits of leopard (Panthera pardus fusca), dhole (Cuon alpinus) and striped hyena (Hyaena hyaena) in a tropical dry thorn forest of southern India. Journal of the Bombay Natural History Society, 104(2), 178–187.

Aronson, M. F., Lepczyk, C. A., Evans, K. L., Goddard, M. A., Lerman, S. B., MacIvor, J. S., … & Vargo, T. (2017). Biodiversity in the city: key challenges for urban green space management. Frontiers in Ecology and the Environment, 15(4), 189–196.

Ashish, K., Ramesh, T., Kalle, R., & Giordano, A. J. (2024). Spatial heterogeneity facilitates coexistence between striped hyaenas and sympatric carnivores. Journal of Biogeography, 51(2), 173–184.

Athreya, V., Odden, M., Linnell, J. D., Krishnaswamy, J., & Karanth, U. (2013). Big cats in our backyards: persistence of large carnivores in a human dominated landscape in India. PloS one, 8(3), e57872.

Athreya, V., Odden, M., Linnell, J. D., Krishnaswamy, J., & Karanth, K. U. (2016). A cat among the dogs: leopard Panthera pardus diet in a human-dominated landscape in western Maharashtra, India. Oryx, 50(1), 156–162.

Baker, P. J., & Harris, S. (2004). The behavioural ecology of red foxes in urban Bristol in Biology and conservation of wild canids (eds. Macdonald, DW & Sillero-Zubiri, C.) 207–216.

Baker, P.J., Funk, S.M., Harris, S. & White, P.C.L. (2000). Flexible spatial organization of urban foxes, Vulpes vulpes, before and during an outbreak of sarcoptic mange. Anim. Behav. 59, 127–146.

Balme, G. A., Hunter, L., & Braczkowski, A. R. (2012). Applicability of age-based hunting regulations for African leopards. PloS one, 7(4), e35209.

Barker, K.J., Cole, E., Courtemanch, A., Dewey, S., Gustine, D., Mills, K., Stephenson, J., Wise, B., & Middleton, A.D. (2023). Large carnivores avoid humans while prioritizing prey acquisition in anthropogenic areas. J. Anim. Ecol., 92, 889–900. 10.1111/1365–2656.13900

Bateman, P. W., & Fleming, P. A. (2012). Big city life: carnivores in urban environments. Journal of Zoology, 287(1), 1–23.

Bhandari, S., Morley, C., Aryal, A., & Shrestha, U.B. (2020). The diet of the striped hyena in Nepal’s lowland regions. Ecol. Evol. 10(15), 7953–7962.

Borchers, D. L., & Efford, M. G. (2008). Spatially explicit maximum likelihood methods for capture–recapture studies. Biometrics, 64(2), 377–385.

Braczkowski, A. R., O’Bryan, C. J., Stringer, M. J., Watson, J. E. M., Possingham, H. P., & Beyer, H. L. (2018). Leopards provide public health benefits in Mumbai, India. Frontiers in Ecology and the Environment.

Carbone, C., & Gittleman, J. L. (2002). A common rule for the scaling of carnivore density. Science, 295(5563), 2273–2276.

Champion, H. G., & Seth, S. K. (1968). A revised survey of the forest types of India. Manager of publications, India.

Chand, M., Yousuf, P., S, F. K., Maletha, S. K., Nautiyal, R., Habib, B., … & Adhikari, B. S. (2025). Wildlife at the urban fringe: patterns of mammalian space and diel detection in an urban sanctuary in Aravalli, India. European Journal of Wildlife Research, 71(5), 117.

Chatterjee, A. B., Sankar, K., Jhala, Y. V., & Qureshi, Q. (2023). Spatio-temporal patterns of co-occurrence of tigers and leopards within a protected area in central India. Web Ecology, 23(1), 17–34.

Chaudhary, P., Kumar, A., Madan, S., Mittal, M., & Dookia, S. (2023). Urban leopards of national capital: estimating density and occupancy of leopard (Panthera pardus) in Asola Bhatti Wildlife Sanctuary, New Delhi, India. Ecol Environ Conserv, 29(10), 383–391.

Chourasia, P., Mondal, K., Sankar, K., & Qureshi, Q. (2012). Food habits of golden jackal (Canis aureus) and striped hyena (Hyaena hyaena) in Sariska Tiger Reserve, Western India. World J. Zool. 7(2), 106–112.

Crooks, K.R. (2002). Relative sensitivities of mammalian carnivores to habitat fragmentation. Conservation Biology, 16, 488–502.

Dahya, M. N., Chaudhary, R., Shah, A., & Kazi, A. (2023). Assemblage, relative abundance and activity pattern of wild mammals in the human-dominated landscape of Vansda taluka, Gujarat, India. Mammalia, 87(5), 434–441.

Davison, J., Huck, M., Delahay, R.J. & Roper, T.J. (2009). Restricted ranging behaviour in a high-density population of urban badgers. J. Zool. (Lond.) 277, 45–53.

Deka, J. R., Lal, U., & Sarma, P. K. (2025). Indian leopards (Panthera pardus fusca) facing space struggles in a Metropolitan district of Northeast India. Environmental Monitoring and Assessment, 197(8), 880.

Desai, B. S. P., D’Costa, A., Praveen Kumar, M.K., & Shyama, S.K. (2021). Diet of Leopards Panthera pardus fusca inhabiting protected areas and human-dominated landscapes in Goa, India. Journal of Threatened Taxa.

Devens, C. H., et al. (2021). Estimating leopard density across the highly modified human-dominated landscape of the Western Cape, South Africa. Oryx 55: 34–45.

Dutta, S. (2024). Study of niche partitioning and interspecific interaction among large carnivores in the dry deciduous ecosystem of Panna Tiger Reserve, Central India. Doctoral dissertation, Zoological Survey of India, Kolkata, Wildlife Institute of India, Dehradun and University of Calcutta, Kolkata, India, 1–202.

Efford, M. (2004). Density estimation in live trapping studies. Oikos, 106(3), 598–610.

Efford, M. G. (2025). The SECR book. A handbook of spatially explicit capture–recapture methods.

Efford, M. G. (2026). secr: Spatially explicit capture–recapture models (Version 5.4.3) [Computer software]. Comprehensive R Archive Network (CRAN). 10.32614/cran.package.secr

Efford, M. G., & Schofield, M. R. (2022). A review of movement models in open population capture–recapture. Methods in Ecology and Evolution, 13(10), 2106–2118.

Esri. (2026). “Imagery” [Basemap]. ArcGIS Online World Imagery. https://www.arcgis.com/home/item.html?id=10df2279f9684e4a9f6a7f08febac2a9

Fedriani, J.M., Fuller, T.K. & Sauvajot, R.M. (2001). Does availability of anthropogenic food enhance densities of omnivorous mammals? An example with coyotes in southern California. Ecography 24, 325–331.

Forest Department, Government of Rajasthan, Forest Department, Government of Rajasthan. https://forest.rajasthan.gov.in/content/raj/forest/en/home.html#. Accessed 22 September 2026.

Forest Survey of India. (2019). India State of Forest Report 2019 (Vol. I & II). Ministry of Environment, Forest and Climate Change, Government of India. https://www.fsi.nic.in/forest-report-2019.

Frid, A., & Dill, L. M. (2002). Human-caused disturbance stimuli as a form of predation risk. Conservation Ecology, 6(1), 11.

Gaynor, K. M., Hojnowski, C. E., Carter, N. H., & Brashares, J. S. (2018). The influence of human disturbance on wildlife nocturnality. Science, 360(6394), 1232–1235.

Gehrt, S. D., Brown, J. L., & Anchor, C. (2011). Is the Urban Coyote a Misanthropic Synanthrope? The Case from Chicago. Cities and the Environment (CATE), 4(1), Article 3.

Gese, E.M. & Bekoff, M. (2004). Coyote Canis latrans Say, 1823. In Canids: foxes, wolves, jackals and dogs. Status survey and conservation action plan, 81–87.

C. Sillero-Zubiri, M. Hoffmann, D.W. Macdonald & IUCN/SSC Canid Specialist Group (Eds). Cambridge: IUCN Publications Services Unit.

Goyal, S. P., Chauhan, D. S., Agrawal, M. K., & Thapa, R. (2000). A study on distribution, relative abundance and food habits of leopard (Panthera pardus) in Garhwal Himalayas. Technical report. Wildlife Institute of India, Dehradun.

Grimm, N.B., Faeth, S.H., Golubiewski, N.E., Redman, C.L., Wu, J., Bai, X. et al. (2008). Global change and the ecology of cities. Science, 319, 756–760.

Groffman, P. M., Cavender-Bares, J., Bettez, N. D., Morgan Grove, J., Hall, S. J., Heffernan, J. B., Hobbie, S. E., Larson, K. L., Morse, J. L., Neill, C., Nelson, K., O’Neil-Dunne, J., Ogden, L., Pataki, D. E., Polsky, C., Chowdhury, R. R., & Steele, M. K. (2014). Ecological homogenization of urban USA. Frontiers in Ecology and the Environment, 12, 74–81.

Gupta, S., Mondal, K., Sankar, K., & Qureshi, Q. (2009). Estimation of striped hyena Hyaena hyaena population using camera traps in Sariska Tiger Reserve, Rajasthan, India. Journal of the Bombay Natural History Society, 106(3), 284.

Gupta, S., Sanyal, A., Saha, G. K., & Ghosh, A. K. (2016). Diurnal activity pattern of Golden Jackal (Canis aureus Linn.) in an urban landscape of Kolkata, India. Proceedings of the Zoological Society, 69(1), 75–80.

Hamer, A. J. & McDonnell, M. J. (2010). The response of herpetofauna to urbanization: inferring patterns of persistence from wildlife databases. Austral Ecology, 35, 568–580.

Hayward, M. W., & Marlow, N. (2014). Will dingoes really conserve wildlife and can our methods tell?. Journal of Applied Ecology, 51(4), 835–838.

Hayward, M. W., Bugir, C., Lyngdoh, S., & Habib, B. (2022). Prey Preferences of the Large Predators of Asia. Case Studies of Wildlife Ecology and Conservation in India, 11-17.

Hayward, M. W., Hayward, G. J., Druce, D. J., & Kerley, G. I. (2009). Do fences constrain predator movements on an evolutionary scale? Home range, food intake and movement patterns of large predators reintroduced to Addo Elephant National Park, South Africa. Biodiversity and Conservation, 18(4), 887–904.

Hayward, M. W., Henschel, P., O’Brien, J., Hofmeyr, M., Balme, G., & Kerley, G. I. (2006). Prey preferences of the leopard (Panthera pardus). Journal of Zoology, 270(2), 298–313.

Hiby, L., Lovell, P., Patil, N., Kumar, N. S., Gopalaswamy, A. M., & Karanth, K. U. (2009). A tiger cannot change its stripes: using a three-dimensional model to match images of living tigers and tiger skins. Biology letters, 5(3), 383.

Hiby, L.R., Reece, J.F., Wright, R., Jaisinghani, R., Singh, B. & Hiby, E.F. (2011). A mark resight survey method to estimate the roaming dog population in three cities in Rajasthan, India. BMC Veterinary Research, 7, 46.

Ives, C. D., Lentini, P. E., Threlfall, C. G., Ikin, K., Shanahan, D. F., Garrard, G. E., … & Kendal, D. (2016). Cities are hotspots for threatened species. Global Ecology and biogeography, 25(1), 117–126.

Jacobson, A. P. et al. (2016). Leopard (Panthera pardus) status, distribution, and the research efforts across its range. PeerJ, 4, 1–28.

Jaipur Municipal Corporation. Nagar Nigam Jaipur. Accessed August 24, 2026, from https://www.jaipurmc.org

Jaipur Development Authority (JDA). (2025). Master Development Plan 2025: Volume 1—Existing Profile Jaipur Region. Jaipur Development Authority, Government of Rajasthan, Jaipur.

Jawaid, M. F., Sharma, M., Pipralia, S., & Kumar, A. (2017). City profile: Jaipur. Cities, 68, 63–81.

Jhala, Y. V., Qureshi, Q., Gopal, R., & Sinha, P. R. (2011). Status of the Tigers, Co-predators, and Prey in India, 2010. National Tiger Conservation Authority, Govt. of India, New Delhi, and Wildlife Institute of India, Dehradun. Dehradun. TR 2011/003 pp-302.

Jhala, Y.V., Qureshi, Q., & Yadav, S.P. (2021). Status of leopards, co-predators, and megaherbivores in India, 2018. National Tiger Conservation Authority, Government of India, New Delhi, and Wildlife Institute of India, Dehradun. ISBN 81-85496-56-0.

Kalle, R., Ramesh, T., Qureshi, Q., & Sankar, K. (2011). Density of tiger and leopard in a tropical deciduous forest of Mudumalai Tiger Reserve, southern India, as estimated using photographic capture–recapture sampling. Acta Theriologica, 56(4), 335–342.

Karanth, K. U., & Nichols, J. D. (1998). Estimation of tiger densities in India using photographic captures and recaptures. Ecology, 79(8), 2852–2862.

Karanth, K. U., & Nichols, J. D. (Eds.). (2002). Monitoring tigers and their prey: A manual for researchers, managers, and conservationists in tropical Asia. Centre for Wildlife Studies, Bangalore, India.

Karanth, K. U., Nichols, J. D., Kumar, N. S., Link, W. A., & Hines, J. E. (2004). Tigers and their prey: predicting carnivore densities from prey abundance. Proceedings of the National Academy of Sciences, 101(14), 4854–4858.

Karanth, K. U., Srivathsa, A., Vasudev, D., Puri, M., Parameshwaran, R., & Kumar, N. S. (2017). Spatio-temporal interactions facilitate large carnivore sympatry across a resource gradient. Proceedings of the Royal Society B: Biological Sciences, 284(1848), 20161860.

Kolekar, A., Hockings, K., Metcalfe, K., & Gubbi, S. (2024). Identifying Priority Areas for the Indian Leopard (Panthera pardus fusca) Within a Shared Landscape. Ecology and Evolution, 14(10), e70404.

Kumar, N., Gupta, U., Malhotra, H., Jhala, Y. V., Qureshi, Q., Gosler, A. G., & Sergio, F. (2019). The population density of an urban raptor is inextricably tied to human cultural practices. Proceedings of the Royal Society B: Biological Sciences, 286(1900), 20182932.

Kumbhojkar, S., Yosef, R., Kosicki, J. Z., Kwiatkowska, P. K., & Tryjanowski, P. (2021). Dependence of the leopard Panthera pardus fusca in Jaipur, India, on domestic animals. Oryx, 55(5), 692–698.

Kumbhojkar, S., Yosef, R., Mehta, A., & Rakholia, S. (2020). A camera-trap home-range analysis of the Indian leopard (Panthera pardus fusca) in Jaipur, India. Animals, 10(9), 1600.

Kumbhojkar, S., Yosef, R., Mehta, A., & Rakholia, S. (2025). Ecosystem Services Provided by Leopards (Panthera pardus fusca) to the Conurbation of Jaipur, India. In: Angeoletto, F., Tryjanowski, P., Fellowes, M.D.E. (eds) Ecology of Tropical Cities, Volume II. Springer, Cham.

Landy, F., Rodary, E., & Calas, B. (2018). Why did leopards kill humans in Mumbai but not in Nairobi? Wildlife management in and around urban national parks. In From urban national parks to natured cities in the global south: The quest for naturbanity (pp. 157–179). Singapore: Springer Singapore.

Lanszki, J., Hayward, M. W., & Nagyapáti, N. (2018). Feeding responses of the golden jackal after reduction of anthropogenic food subsidies. PLoS One, 13(12), e0208727.

Latafat, K., Sadhu, A., Qureshi, Q., & Jhala, Y. V. (2023). Abundance and activity of carnivores in two protected areas of semi-arid western India with varying top predator density and human impacts. European Journal of Wildlife Research, 69(1), 15.

Leighton, G. R., Bishop, J. M., Merondun, J., Winterton, D. J., O’Riain, M. J., & Serieys, L. E. (2022). Hiding in plain sight: risk mitigation by a cryptic carnivore foraging at the urban edge. Animal Conservation, 25(2), 244–258.

Lindstedt, S. L., B. J. Miller, and S. W. Buskirk. 1986. Home range, time, and body size in mammals. Ecology 67:413–418.

Macdonald, D. W. (1983). The ecology of carnivore social behaviour. Nature, 301(5899), 379–384.

Majgaonkar, I., Vaidyanathan, S., Srivathsa, A., Shivakumar, S., Limaye, S., & Athreya, V. (2019). Land-sharing potential of large carnivores in human-modified landscapes of western India. Conservation Science and Practice, 1(5), e34.

Mandal, D. K. (2018). Ecology of striped hyena (Hyaena hyaena) in Sariska Tiger Reserve, Rajasthan. Doctoral dissertation, PhD Thesis, Saurashtra University, Rajkot, India.

Mandal, D., Chatterjee, D., Qureshi, Q., & Sankar, K. (2018). Behavioural observations on interaction of leopard and striped hyena, western India. Cat News, 67, 20–21.

McKinney, M. L. (2006). Urbanization as a major cause of biotic homogenization. Biological Conservation, 127(3), 247–260.

McNab, B. K. 1963. Bioenergetics and the determination of home range. The American Naturalist 97:133–140.

Meredith, M., & Ridout, M. (2014). Overview of the overlap package. R. Proj, 1(9).

Meredith, M., Ridout, M., & Campbell, L. A. (2013). Overlap: estimates of coefficient of overlapping for animal activity patterns. CRAN: Contributed Packages.

Miller, J. R., Pitman, R. T., Mann, G. K., Fuller, A. K., & Balme, G. A. (2018). Lions and leopards coexist without spatial, temporal or demographic effects of interspecific competition. Journal of Animal Ecology, 87(6), 1709–1726.

Monterroso, P., Alves, P. C., & Ferreras, P. (2014). Plasticity in circadian activity patterns of mesocarnivores in Southwestern Europe: implications for species coexistence. Behavioral Ecology and Sociobiology, 68(9), 1403–1417.

Newsome, T. M., Dellinger, J. A., Pavey, C. R., Ripple, W. J., Shores, C. R., Wirsing, A. J., & Dickman, C. R. (2015). The ecological effects of providing resource subsidies to predators. Global Ecology and Biogeography, 24(1), 1–11.

O’Brien, T. G., Kinnaird, M. F., & Wibisono, H. T. (2003, May). Crouching tigers, hidden prey: Sumatran tiger and prey populations in a tropical forest landscape. In Animal Conservation Forum (Vol. 6, No. 2, pp. 131–139). Cambridge University Press.

Odden, M., Athreya, V., Rattan, S., & Linnell, J. D. (2014). Adaptable neighbours: movement patterns of GPS-collared leopards in human dominated landscapes in India. PloS one, 9(11), e112044.

Oke, T.R. (1982). The energetic basis of the urban heat island. Q. J. R. Meteorol. Soc., 108, 1–24.

Oriol-Cotterill, A., Valeix, M., Frank, L. G., Riginos, C., & Macdonald, D. W. (2015). Landscapes of coexistence for terrestrial carnivores: The ecological consequences of being downgraded from ultimate to penultimate predator by humans. Oikos, 124(10), 1263–1273.

Panda, D., Krishna, J. S., Sharma, A., Pandey, P., Lee, H., & Singh, R. (2023). Ecosystem services provided by striped hyenas in the human-dominated landscape of Rajasthan, India. Science of The Total Environment, 903, 166267.

Panda, D., Mohanty, S., Suryan, T., Pandey, P., Lee, H., & Singh, R. (2022). High striped hyena density suggests coexistence with humans in an agricultural landscape, Rajasthan. PloS one, 17(5), e0266832.

Patra, M. K., Jain, D., Sadhu, A., Qureshi, Q., & Jhala, Y. V. (2026). Spatio-temporal niche partitioning and interactions between leopards and striped hyenas under anthropogenic disturbance in the Indian Terai Arc Landscape. Mammal Research, 71(4), 63.

Périquet, S., Fritz, H., & Revilla, E. (2015). The Lion King and the Hyaena Queen: large carnivore interactions and coexistence. Biological Reviews, 90(4), 1197–1214.

Qureshi, Q., Jhala, Y. V., Yadav, S. P., & Mallick, A. (2023). Status of tigers, co-predators and prey in India, 2022. National Tiger Conservation Authority, Government of India, New Delhi, and Wildlife Institute of India, Dehradun, 3, 50–56.

R Core Team. (2026). R: A language and environment for statistical computing (Version 4.6. 0)[Computer software]. R Foundation for Statistical Computing.

Reshamwala, H. S., Shrotriya, S., Bora, B., Lyngdoh, S., Dirzo, R., & Habib, B. (2018). Anthropogenic food subsidies change the pattern of red fox diet and occurrence across Trans-Himalayas, India. Journal of Arid Environments, 150, 15–20.

Ridout, M. S., & Linkie, M. (2009). Estimating overlap of daily activity patterns from camera trap data. Journal of Agricultural, Biological, and Environmental Statistics, 14(3), 322–337.

Rowcliffe, M., & Rowcliffe, M. M. (2016). Package ‘activity’. Animal activity statistics R package version, 1(10.32614).

Šálek, M., Drahníková, L., & Tkadlec, E. (2015). Changes in home range sizes and population densities of carnivore species along the natural to urban habitat gradient. Mammal Review, 45(1), 1–14.

Savard, J. P. L., Clergeau, P., & Mennechez, G. (2000). Biodiversity concepts and urban ecosystems. Landscape and Urban Planning, 48, 131–142.

Scott, D. M., Berg, M. J., Tolhurst, B. A., Chauvenet, A. L., Smith, G. C., Neaves, K., … & Baker, P. J. (2014). Changes in the distribution of red foxes (Vulpes vulpes) in urban areas in Great Britain: findings and limitations of a media-driven nationwide survey. PloS one, 9(6), e99059.

Searle, C. E., Smit, J. B., Cusack, J. J., Strampelli, P., Grau, A., Mkuburo, L., … & Dickman, A. J. (2021). Temporal partitioning and spatiotemporal avoidance among large carnivores in a human-impacted African landscape. PLoS One, 16(9), e0256876.

Sehgal, J. J., Panda, D., Kumar, D., Kalsi, R., Allen, M. L., & Singh, R. (2023). Does the size of a protected area matter? An assessment of leopard population and habitat usage in a protected area of Shiwalik foothills, Himalaya. Mammal Research, 68(4), 459–469.

Seto, K.C., Güneralp, B., & Hutyra, L.R. (2012). Global forecasts of urban expansion to 2030 and direct impacts on biodiversity and carbon pools. P Natl Acad Sci USA, 109, 16083–88.

Shah, G. M., Jan, U., Bhat, B. A., Ahmad, F., & Ahmad, J. (2009). Food habits of the Leopard Panthera pardus in Dachigam National Park, Kashmir, India. Journal of Threatened Taxa, 1(3), 184.

Sharma, N., Gaur, S., Dhyani, R., & Singh, A. (2016). Challenges of small protected areas in urban cities: a case study of Okhla Bird Sanctuary, India. Environment, Development and Sustainability, 18(1), 295–310.

Singh, H. S. (2005). Status of the leopard Panthera pardus fusca in India. Cat News, 42.

Smith, J. A., Suraci, J. P., Clinchy, M., Crawford, A., Roberts, D., Zanette, L. Y., & Wilmers, C. C. (2017). Fear of the human ‘super predator’ reduces feeding time in large carnivores. Proceedings of the Royal Society B: Biological Sciences, 284(1857), 20170433.

Soanes, K., & Lentini, P. E. (2019). When cities are the last chance for saving species. Frontiers in Ecology and the Environment, 17(4), 225–231.

Soulsbury, C. D., & White, P. C. (2015). Human–wildlife interactions in urban areas: a review of conflicts, benefits and opportunities. Wildlife research, 42(7), 541–553.

Surve, N. (2015). Ecology of leopard in Sanjay Gandhi National Park, Maharashtra with special reference to its abundance, prey selection and food habit. Mumbai: Wildlife Institute of India; Maharashtra Forest Department, SGNP.

Surve, N.S., Sathyakumar, S., Sankar, K., Jathanna, D., Gupta, V., & Athreya, V. (2022). Leopards in the City: The Tale of Sanjay Gandhi National Park and Tungareshwar Wildlife Sanctuary, Two Protected Areas in and Adjacent to Mumbai, India. Front. Conserv. Sci. 3, 787031.

Tanwar, K. S., Sadhu, A., & Jhala, Y. V. (2021). Camera trap placement for evaluating species richness, abundance, and activity. Scientific Reports, 11, 23050.

Terborgh, J. (1974). Preservation of natural diversity: the problem of extinction-prone species. Bioscience, 24, 715–722.

Tiwari, S., Karnatak, R. & Goyal, S. P. (2008). Food habits of leopard (Panthera pardus) in Sariska Tiger Reserve, Rajasthan, India. Journal of the Bombay Natural History Society, 105(2), 194–198.

United Nations, Department of Economic and Social Affairs, Population Division (2019). World Urbanization Prospects: The 2018 Revision (ST/ESA/SER.A/420). New York: United Nations.

Vijayan, S., & Pati, B. P. (2002). Impact of changing cropping patterns on man-animal conflicts around Gir Protected Area with specific reference to Talala Sub-District, Gujarat, India. Population and Environment, 23(6), 541–559.

Vissia, S., & Van Langevelde, F. (2022). The effect of body size on co-occurrence patterns within an African carnivore guild. Wildlife Biology, 2022(2), e01004.

Wattegedera, M., Silva, D., Sooriyabandara, C., Wimaladasa, P., Siriwardena, R., Piyasena, M., … & Mahela, M. (2022). A multi-point identification approach for the recognition of individual leopards (Panthera pardus kotiya). Animals, 12(5), 660.

Webbon, C.C., Baker, P.J. & Harris, S. (2004). Faecal density counts for monitoring changes in red fox numbers in rural Britain. J. Appl. Ecol. 41, 768–779.

Woodroffe, R. (2000). Predators and people: using human densities to interpret declines of large carnivores. Animal Conservation, 3, 165–173.

Yirga, G., Ersino, W., De Iongh, H. H., Leirs, H., Gebrehiwot, K., Deckers, J., & Bauer, H. (2013). Spotted hyena (Crocuta crocuta) coexisting at high density with people in Wukro district, northern Ethiopia. Mammalian Biology, 78(3), 193–197.

Yosef, R., & Kumbhojkar, S. (2025). Kleptoparasitism and Coexistence: Resource competition between Indian leopards and striped hyenas. Animals, 15(6), 784.

